# Two major epidemics of highly pathogenic avian influenza virus H5N8 and H5N1 in domestic poultry in France, 2020-2022

**DOI:** 10.1101/2022.06.20.496805

**Authors:** Sébastien Lambert, Benoit Durand, Mathieu Andraud, Roxane Delacourt, Axelle Scoizec, Sophie Le Bouquin, Séverine Rautureau, Billy Bauzile, Claire Guinat, Lisa Fourtune, Jean-Luc Guérin, Mathilde C. Paul, Timothée Vergne

**Affiliations:** IHAP, Université de Toulouse, INRAE, ENVT, Toulouse, France; Agence Nationale de Sécurité Sanitaire de l’Alimentation, de l’Environnement et du Travail, Université Paris-Est, Maisons-Alfort, France; Agence Nationale de Sécurité Sanitaire de l’Alimentation, de l’Environnement et du Travail, Ploufragan, France; Direction Générale de l’Alimentation, Paris, Fance; Department of Biosystems Science and Engineering, ETH Zurich, Basel, Switzerland; Swiss Institute of Bioinformatics, Lausanne, Switzerland

**Keywords:** Influenza A Virus, H5N1 Subtype, Influenza A Virus, H5N8 Subtype, Disease Outbreaks, Incidence, Basic Reproduction Number, Area Under Curve

## Abstract

The spread of highly pathogenic avian influenza (HPAI) viruses worldwide has serious consequences for animal health and a major economic impact on the poultry production sector. Since 2014, Europe has been severely hit by several HPAI epidemics, with France being the most affected country. Most recently, France was again affected by two devastating highly pathogenic avian influenza epidemics in 2020-21 and 2021-22. We conducted a descriptive analysis of the 2020-21 and 2021-22 epidemics, in a first step towards identifying the poultry sector’s remaining vulnerabilities regarding HPAI viruses in France. We examined the spatio-temporal distribution of outbreaks that occurred in France in 2020-21 and 2021-22, and we assessed the outbreaks’ spatial distribution in relation to two High-Risk Zones recently incorporated into French legislation to strengthen HPAI prevention and control. There were 468 reported outbreaks during the 2020-21 epidemic, and 1,223 outbreaks during the 2021-22 epidemic. In both epidemics, most outbreaks (80.6% and 74.0%) were located into the two High-Risk Zones. The southwestern High-Risk Zone was affected in both epidemics, while the western High-Risk Zone was affected for the first time in 2021-22, explaining the extremely high number of outbreaks reported. We showed that the spatial distribution model used to create the two High-Risk Zones was able to predict the location of outbreaks for the 2020-21 and 2021-22 epidemics. These zones were characterized by high poultry farm densities; future efforts should therefore focus on reducing the density of susceptible poultry in highly dense areas.

## Introduction

Unprecedented spread of highly pathogenic avian influenza (HPAI) viruses was observed across Europe, Asia, Africa and North America in the winter of 2021-22, infecting tens of millions of poultry birds and hundreds of thousands of wild birds (Miller, 2022; Wille and Barr, 2022). These viruses may cause severe clinical signs and high mortality rates in birds, causing serious economic losses in poultry and raising issues for the conservation of vulnerable wild bird species (Miller, 2022; Wille and Barr, 2022). HPAI viruses are also concerning for public health because of their zoonotic potential and the risk of spillover to people, which increases as the number of bird outbreaks increases (Miller, 2022; Wille and Barr, 2022).

In Europe, several HPAI subtype H5Nx epidemic waves occurred since the emergence of the current circulating lineage 2.3.4.4 and its introduction by wild migratory birds in late 2014. In particular, during the winter of 2016-17, Europe was affected by an unexpectedly large HPAI subtype H5N8 epidemic, with 1,218 poultry farm outbreaks reported across 29 countries (EFSA et al., 2022). During this epidemic, France was the most heavily affected European country, with 464 poultry farm outbreaks (Guinat et al., 2018). The control measures implemented included culling of infected farms (IFs) and pre-emptive culling of farms around IFs. These measures led to more than 6.8 million poultry being culled and caused a substantial economic impact on the French poultry industry (Guinat et al., 2018). Several studies highlighted the role of biosecurity practices, poultry farm density, and duck transportation in the spread of HPAI viruses between French farms during the 2016-17 epidemic (Guinat et al., 2019; Guinat, Comin et al., 2020; Guinat, Durand et al., 2020; Bauzile, Sicard et al., 2022).

Following the 2015-16 and 2016-17 HPAI epidemics, new regulations on biosecurity for poultry farms and live bird transportation came into effect in France (DGAl, 2016, 2018). A national training program was made mandatory for poultry producers, and the French veterinary authorities performed biosecurity audits on poultry farms between 2016 and 2018 to verify compliance with the new regulations (Delpont et al., 2021). Two “High-Risk Zones” (HRZ) were also incorporated into French legislation in September 2021 (DGAl, 2021), where pre-emptive measures (such as confining ducks indoors and timely pre-movement testing) were implemented in autumn and winter when the risk of HPAI introduction increased. These two HRZ, located in the southwestern and western parts of France, were created based on a spatial distribution model of the 2016-17 outbreaks, and were characterized by high poultry farm densities and high duck movement numbers (Guinat et al., 2019).

In the winters of 2020-21 and 2021-22, France and Europe were affected by two other major HPAI epidemics (subtypes H5N8 and H5N1, respectively), with the number of poultry farm outbreaks exceeding those caused by the 2016-17 epidemic. Given the improved biosecurity and drastic control measures implemented in France, these latest epidemics are worrying. Our study therefore aimed to characterize the spatio-temporal patterns of the 2020-21 and 2021-22 epidemics in France and to assess the outbreaks’ spatial distribution in relation to the two HRZ. This constitutes a first step towards identifying the poultry sector’s remaining vulnerabilities regarding HPAI viruses in France.

## Materials and Methods

### Data collection

We obtained data on the HPAI poultry farm outbreaks for the 2020-21 (December 2, 2020-March 20, 2021) and 2021-22 (November 21, 2021-April 12, 2022) epidemics from the French General Directorate for Food (DGAl) of the French Ministry of Agriculture. An outbreak was defined as detection of at least one laboratory confirmed HPAI-infected bird (by virus isolation or polymerase chain reaction) in a commercial poultry farm. Data included the species involved, type of production, date of suspicion (by clinical or active surveillance) and geographical location of each outbreak. Spatial data were obtained from DGAl for the HRZ, and from Guinat et al. (Guinat et al., 2019) for the predicted probability of having at least one HPAI outbreak in a commune (the smallest administrative unit in France, corresponding to Nomenclature of Territorial Units for Statistics level 5).

### Descriptive analysis

All analyses were conducted using R statistical software version 4.1.1 (R Core Team, 2021). Epidemic curves were plotted using the R package incidence (Kamvar et al., 2019; Jombart et al., 2020). All maps were produced using the R package tmap (Tennekes, 2018). Geographic data of all countries and administrative areas were downloaded from the GADM (https://gadm.org/) database using the R package raster (Hijmans, 2021). Outbreaks for which the precise location was missing were given the coordinates of the centroid of the commune where they occurred. The coordinates of the communes’ centroids were obtained from the French National Institute of Geographic and Forest Information (IGN) ADMIN EXPRESS database (https://geoservices.ign.fr/adminexpress). To assess the ability of the 2016-17 model (Guinat et al., 2019) to predict the location of outbreaks for the 2020-21 and 2021-22 epidemics, we calculated the area under the receiver operator characteristic curves (AUC) for both epidemics using the R package pROC (Robin et al., 2011).

### Transmission dynamics (effective reproduction number)

To study the transmission dynamics, we estimated the effective reproduction number *R*_e_ between farms (i.e., the average number of secondary farms infected by each infectious farm) using the approach of Wallinga and Teunis (Wallinga and Teunis, 2004). The approach of Wallinga and Teunis allows estimating *R*_e_ based on the time variations of incidence and on the distribution of the serial interval (time interval between symptom onset in a farm and that of its secondary cases). Because the distribution of the serial interval distribution was unknown, we first used the approach of White and Pagano (White and Pagano, 2008) to estimate simultaneously the basic reproduction number *R*_0_ and the mean and standard deviation of the discretized serial interval distribution (assumed to follow a gamma distribution), based on the initial exponential phase of the epidemic. This approach was implemented using the R package R0 (Obadia et al., 2012; Boelle and Obadia, 2015). With the estimates of the serial interval distribution parameters, we were then able to estimate the effective reproduction number using the approach of Wallinga and Teunis (Wallinga and Teunis, 2004), implemented in the R package EpiEstim (Cori, 2021).

The estimates of the approach of White and Pagano (White and Pagano, 2008) implemented in the R package R0 can be sensitive to the selected time period over which epidemic growth is considered exponential (Obadia et al., 2012). By default, the time period considered is from the date of the first case up to the date of the maximum daily incidence (Obadia et al., 2012; Boelle and Obadia, 2015). Another possibility is to select the time period producing the largest R-squared value, corresponding to the period over which the model fitted the data best (Obadia et al., 2012). To assess the sensitivity of the *R*_e_ estimation to the parameters of the serial interval distribution, we used the approach of Wallinga and Teunis (Wallinga and Teunis, 2004) again, this time using the mean and standard deviation of the serial interval distribution that produced the highest R-squared value.

## Results and Discussion

The 2020-21 epidemic consisted of a single wave with 468 outbreaks (Figure 1A) clustered in southwestern France (Figure 2A). In contrast, the 2021-22 epidemic was characterized by 1,223 outbreaks (as of April 12, 2022) divided into two spatio-temporal clusters, the first in the southwest and the second in the western part of the country (Figure 1B and Figure 2B), with a higher incidence during the second wave. Most outbreaks were located in the HRZ, both in 2020-21 (80.6% of the outbreaks) and 2021-22 (74.0% - Figure 1A and B, Figure 2). The AUCs of the 2016-17 model were 0.86 (95% confidence interval: 0.83-0.89) for the 2020-21 epidemic and 0.86 (95%CI: 0.85-0.88) for the 2021-22 epidemic.

**Figure 1:**
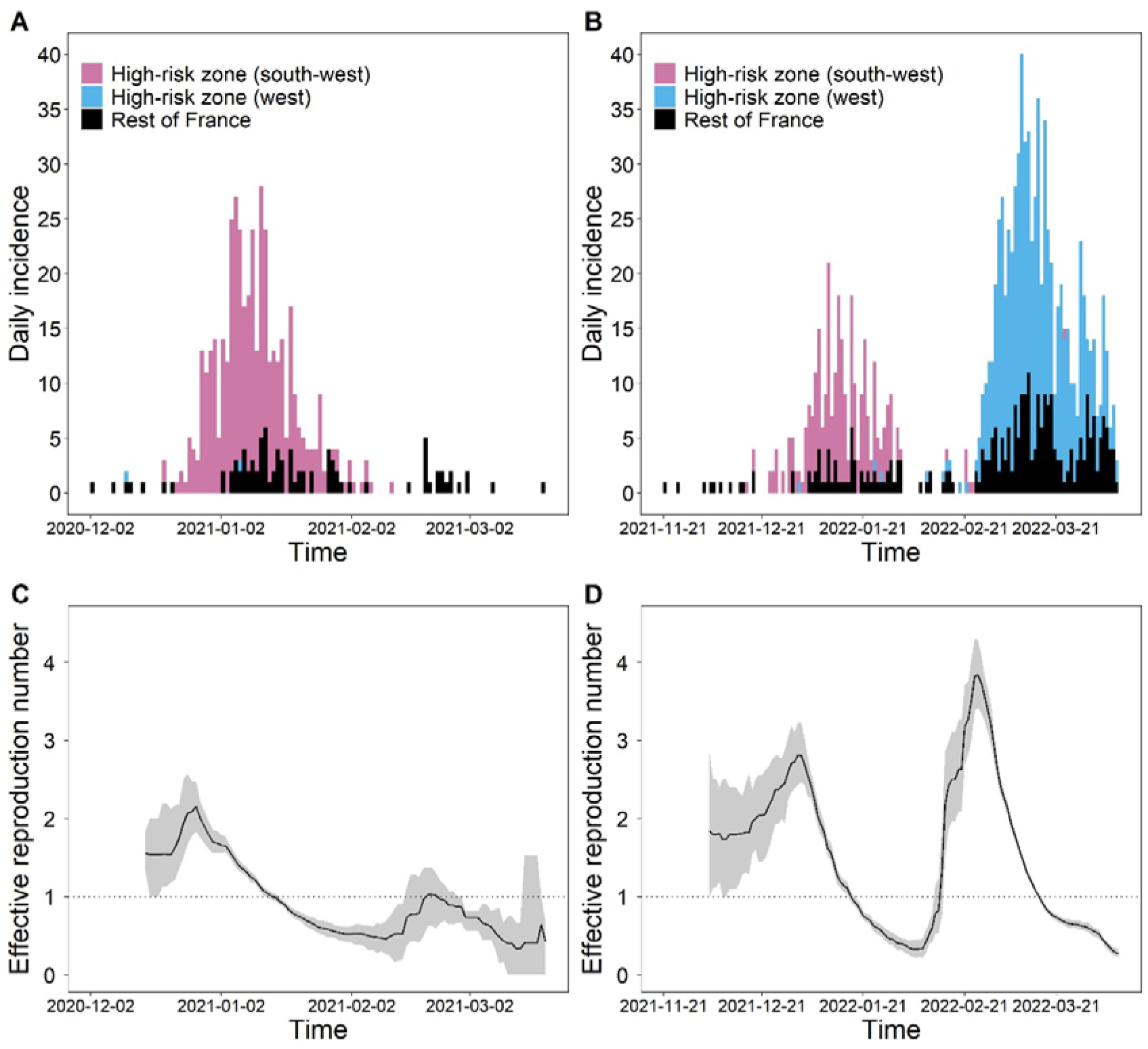
Incidence (A, B) and estimated effective reproduction number (C, D) of the 2020-21 (A, C) and 2021-22 (B, D) HPAI virus epidemics in commercial poultry farms in France. In panels C and D, the graph shows, at each day, the estimate of the effective reproduction number over the 14-day window finishing on that day. The line represents the mean and the grey area represent the 95% confidence interval.

**Figure 2:**
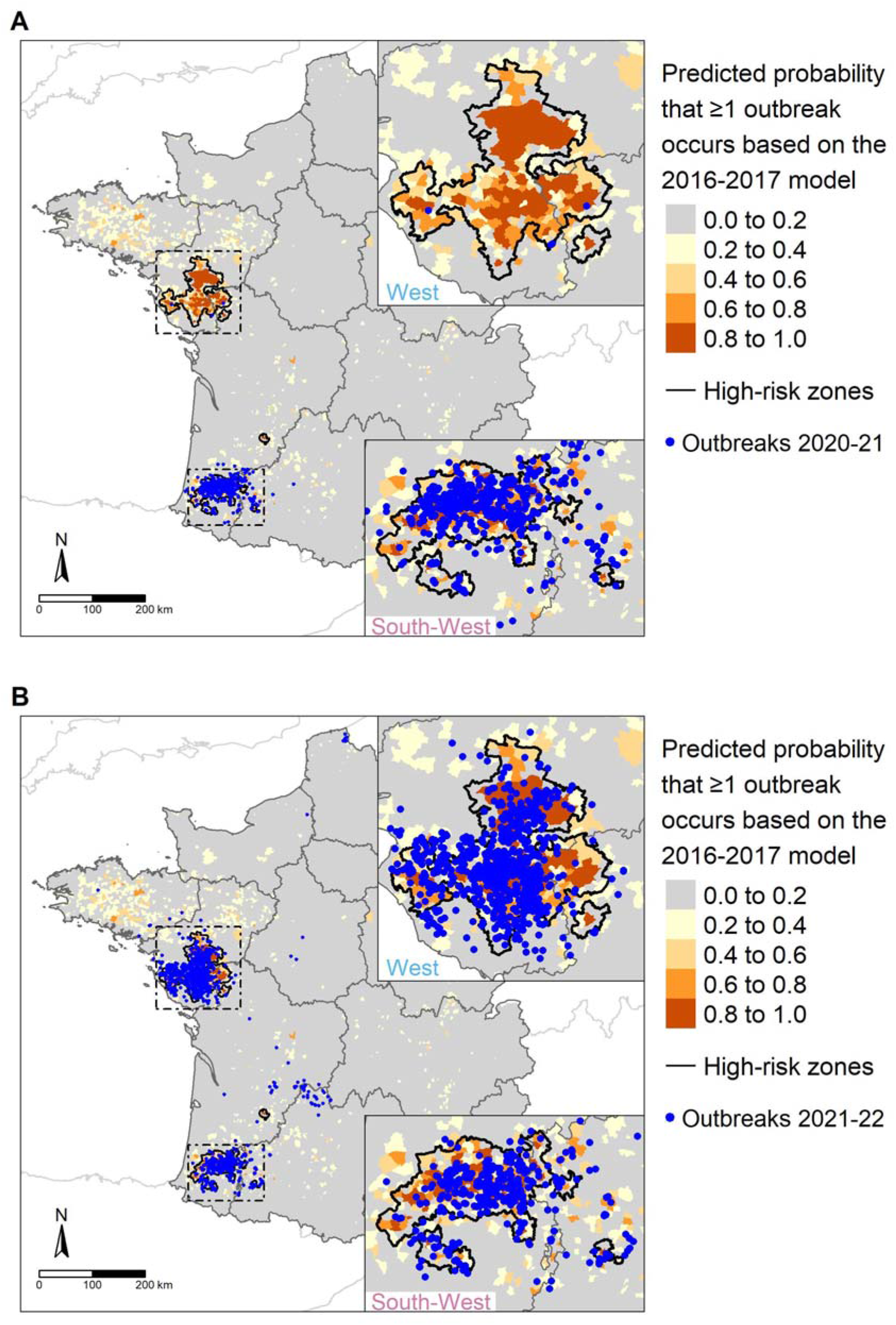
HPAI virus outbreaks in commercial poultry farms in France in 2020-21 (A) and 2021-22 (B), and predicted probability of having at least one outbreak in a commune according to the 2016-2017 model of Guinat *et al*. (2019) (2019).

In both epidemics, the vast majority of farm outbreaks in the southwestern HRZ (82.7% and 69.9%) were reported in ducks (Table 1), mainly in farms raising ducks for *foie gras* production (breeding and/or force-feeding production stages in Table 1). Conversely, in the western HRZ in 2021-22, only half of the outbreaks were reported in duck farms, mainly in breeder and broiler farms (Table 1). Most of the other outbreaks were reported in galliform farms (mainly chickens and turkeys - Table 1).

**Table 1:** Distribution of HPAI virus outbreaks in commercial poultry farms in France during the 2020-21 (H5N8) and 2021-22 (H5N1) epidemics.

|  | 2020-21 |  |  |  | 2021-22 |  |  |  |  |  |
| --- | --- | --- | --- | --- | --- | --- | --- | --- | --- | --- |
|  | South-West<sup>*</sup> |  | TOTAL |  | South-West<sup>*</sup> |  | West<sup>†</sup> |  | TOTAL |  |
| Type of species |  |  |  |  |  |  |  |  |  |  |
| Duck | 310 | (82.7%) | 389 | (83.1%) | 193 | (69.9%) | 315 | (50.1%) | 711 | (58.1%) |
| Goose | 1 | (0.3%) | 1 | (0.2%) | 4 | (1.4%) | 4 | (0.6%) | 8 | (0.7%) |
| Chicken | 34 | (9.1%) | 42 | (9.0%) | 45 | (16.3%) | 172 | (27.3%) | 287 | (23.5%) |
| Turkey | - | - | - | - | - | - | 77 | (12.2%) | 101 | (8.3%) |
| Other galliformes | 6 | (1.6%) | 7 | (1.5%) | 5 | (1.8%) | 35 | (5.6%) | 44 | (3.6%) |
| Multispecies (galliformes) | 2 | (0.5%) | 2 | (0.4%) | 1 | (0.4%) | - | - | 4 | (0.3%) |
| Multispecies (palmipeds + galliformes) | 22 | (5.9%) | 27 | (5.8%) | 5 | (1.8%) | 6 | (1.0%) | 14 | (1.1%) |
| Pigeon | - | - | - | - | - | - | 2 | (0.3%) | 2 | (0.2%) |
| Unknown | - | - | - | - | 23 | (8.3%) | 18 | (2.9%) | 52 | (4.3%) |
| TOTAL | 375 | (100%) | 468 | (100%) | 276 | (100%) | 629 | (100%) | 1,223 | (100%) |
| Type of production for duck farms |  |  |  |  |  |  |  |  |  |  |
| Breeding | 211 | (68.1%) | 260 | (66.8%) | 120 | (62.2%) | 64 | (20.3%) | 276 | (38.8%) |
| Force-feeding | 76 | (24.5%) | 97 | (24.9%) | 52 | (26.9%) | 40 | (12.7%) | 132 | (18.6%) |
| Breeding + force-feeding | 1 | (0.3%) | 1 | (0.3%) | - | - | - | - | - | - |
| Broiler | 2 | (0.6%) | 7 | (1.8%) | 6 | (3.1%) | 126 | (40.0%) | 178 | (25.0%) |
| Breeder | 7 | (2.3%) | 8 | (2.1%) | 1 | (0.5%) | 53 | (16.8%) | 68 | (9.6%) |
| Other | 2 | (0.6%) | 3 | (0.8%) | 2 | (1.0%) | 19 | (6.0%) | 25 | (3.5%) |
| Unknown | 11 | (3.5%) | 13 | (3.3%) | 12 | (6.2%) | 13 | (4.1%) | 32 | (4.5%) |
| TOTAL | 310 | (100%) | 389 | (100%) | 193 | (100%) | 315 | (100%) | 711 | (100%) |
| Type of production for chicken farms |  |  |  |  |  |  |  |  |  |  |
| Broiler | 26 | (76.5%) | 33 | (78.6%) | 32 | (71.1%) | 101 | (58.7%) | 164 | (57.1%) |
| Layer | 4 | (11.8%) | 5 | (11.9%) | 7 | (15.6%) | 32 | (18.6%) | 65 | (22.6%) |
| Breeder | 3 | (8.8%) | 3 | (7.1%) | 1 | (2.2%) | 20 | (11.6%) | 26 | (9.1%) |
| Other | - | - | - | - | - | - | 15 | (8.7%) | 22 | (7.7%) |
| Unknown | 1 | (2.9%) | 1 | (2.4%) | 5 | (11.1%) | 4 | (2.3%) | 10 | (3.5%) |
| TOTAL | 34 | (100%) | 42 | (100%) | 45 | (100%) | 172 | (100%) | 287 | (100%) |

<sup>\*</sup> Southwestern high-risk zone,
<sup>†</sup> Western high-risk zone (note that in 2020-21, only two outbreaks were located in the western high-risk zone, and were therefore not detailed here).

The estimates of the average serial interval were 4.78 days (standard deviation: 4.63 days) for the 2020-21 epidemic, and 8.9 days (standard deviation: 4.08 days) for the 2021-22 epidemic (Supporting Information Figure S1). In the winter of 2020-21, the *R*_e_ peaked at 2.2 (95%CI: 1.7-2.5) in late December 2020, when the incidence increased in the southwestern HRZ, then decreased below 1 from mid-January 2021 (Figure 1C). In 2021-22, the *R*_e_ first peaked at 2.8 (95%CI: 2.5-3.1) in early January 2022, when the virus was mostly circulating in the southwestern HRZ, and then decreased below 1 in the second half of January 2022 (Figure 1D). At the beginning of February 2022, the *R*_e_ increased again dramatically when the virus reached the western HRZ, peaked at 3.8 (95%CI: 3.4-4.3) at the end of February 2022 and then decreased below 1 by mid-March 2022 (Figure 1D).

The temporal variations of the *R*_e_ estimated using the serial interval distribution parameters of White and Pagano’s default model (Figure 1C-D) matched qualitatively and quantitatively with the *R*_e_ dynamics estimated using the serial interval distribution parameters of White and Pagano’s model with the highest R-squared value (Supporting Information Figure S2), although in the latter the *R*_e_ seemed to peak at higher values in 2021-22.

During the 2020-21 and 2021-22 epidemics, the virus circulated extensively in the southwestern HRZ. In that region, where the density of farms raising ducks for *foie gras* production is extremely high, it is worth noting that the number of reported outbreaks decreased from 375 to 276 between the two epidemics, likely the result of a higher level of awareness among farmers, more effective implementation of control strategies and a decrease of duck flock density. However, in the winter of 2021-22, a second spatio-temporal cluster of outbreaks occurred in the western HRZ, with no clear epidemiological link with the southwestern cluster (EFSA et al., 2022). The spread of HPAI within the western HRZ occurred for the first time during the 2021-22 epidemic and explains the extremely high number of outbreaks reported. The species composition in poultry farms in this zone, combined with a higher flock size on average, may have had a significant impact on the pattern of the epidemic in this area. Why the virus spread in the western HRZ in 2021-22 but not in the previous epidemics remains to be determined.

Biosecurity and control measures were significantly improved after 2016. The occurrence of these two major epidemics would suggest that these improvements were not sufficient to prevent the spread of the virus. Interestingly, although the 2016-17 outbreaks only clustered in the southwest part of France, Guinat et al. (Guinat et al., 2019) identified two HRZ that predicted with high accuracy the two spatial clusters observed during the 2021-22 epidemic (Figure 2B). The main risk factors identified were density of poultry farms and activities related to duck movements. The latter risk factor has already been the target of considerable improvement measures. To increase the resilience of the poultry sector in France, and other hardly-hit European countries, future efforts should therefore focus on reducing the density of susceptible poultry farms and the number of susceptible birds on farms in high-density areas during the high risk periods (Bauzile, Durand et al., 2022).

Vaccinating domestic poultry against avian influenza is generally prohibited in the European Union due to the trade restrictions it would generate (European Commission, 2006). However, this once-tabooed prevention strategy is currently being given full consideration in Europe, as it is becoming clear that the accelerating pace of occurrence of devastating HPAI epidemics is generating new challenges that cannot be addressed with more traditional prevention and control approaches alone (EFSA et al., 2021; Stokstad, 2022; Wille and Barr, 2022). In the long term, restructuration of the European poultry sector in densely populated poultry areas, although challenging, may be required alongside vaccination to control HPAI epidemics (EFSA et al., 2021; Stokstad, 2022). Therefore, further research is needed to devise new suitable and sustainable HPAI mitigation strategies in Europe.

## Acknowledgements

This study was performed in the framework of the “Chair for Avian Biosecurity”, hosted by the National Veterinary College of Toulouse and funded by the Direction Générale de l’Alimentation, Ministère de l’Agriculture et de l’Alimentation, France. Claire Guinat is funded by the European Union’s Horizon 2020 research and innovation program under the Marie Sklodowska-Curie grant agreement No 842621. The funders had no role in the design of the study; in the collection, analyses, or interpretation of data; in the writing of the manuscript, or in the decision to publish the results. The authors wish to thank Grace Delobel for language editing.

