## Supporting Information for "Two major epidemics of highly pathogenic avian influenza virus H5N8 and H5N1 in domestic poultry in France, 2020-2022"

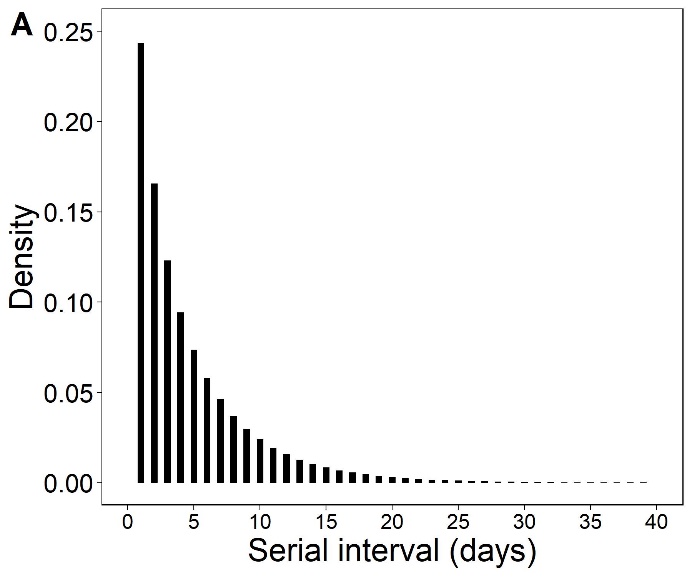

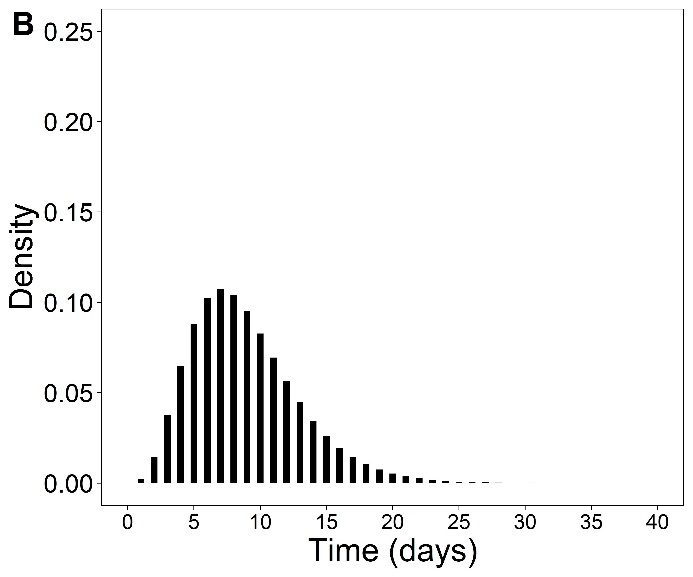

**Figure S1. Discrete distributions of the serial interval of highly pathogenic avian influenza in poultry farms in France. A: 2020-21 (H5N8), mean 4.78 days and standard deviation 4.63 days; B: 2021-22 (H5N1), mean 8.9 days and standard deviation 4.08 days.**

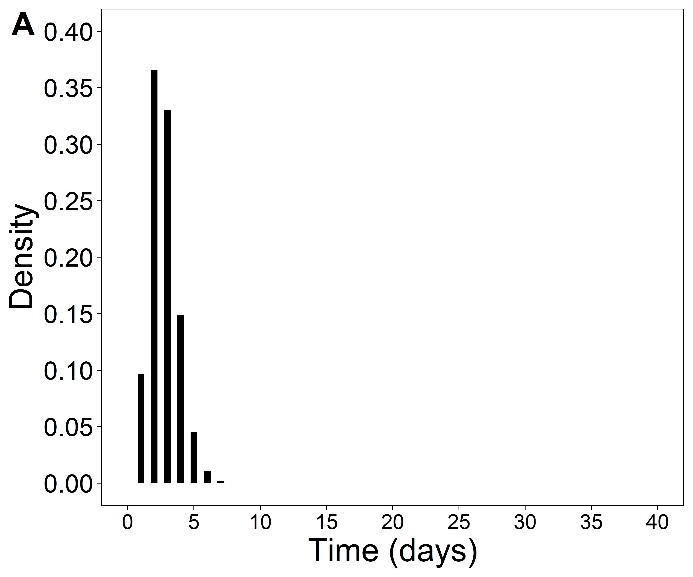

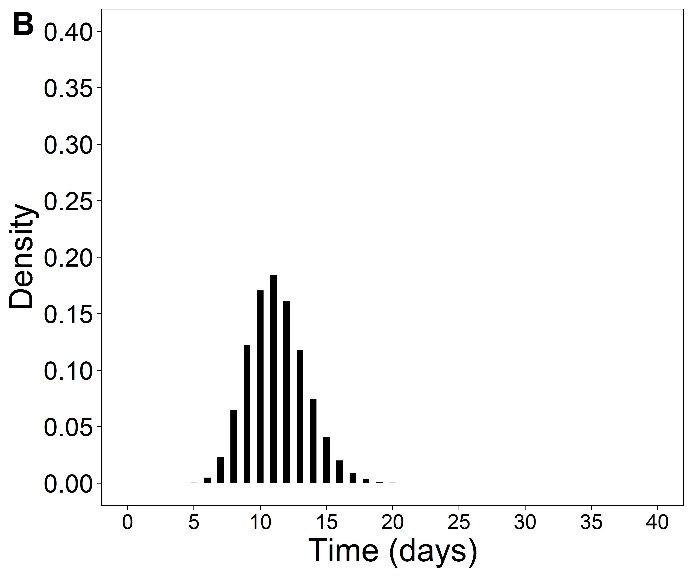

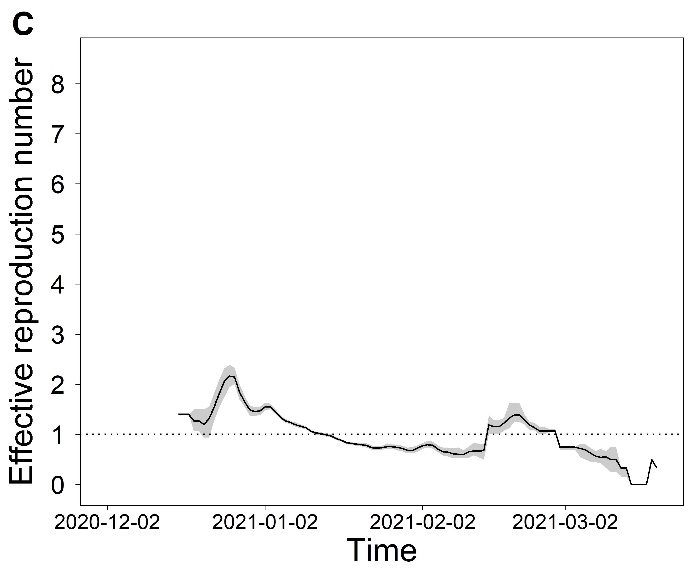

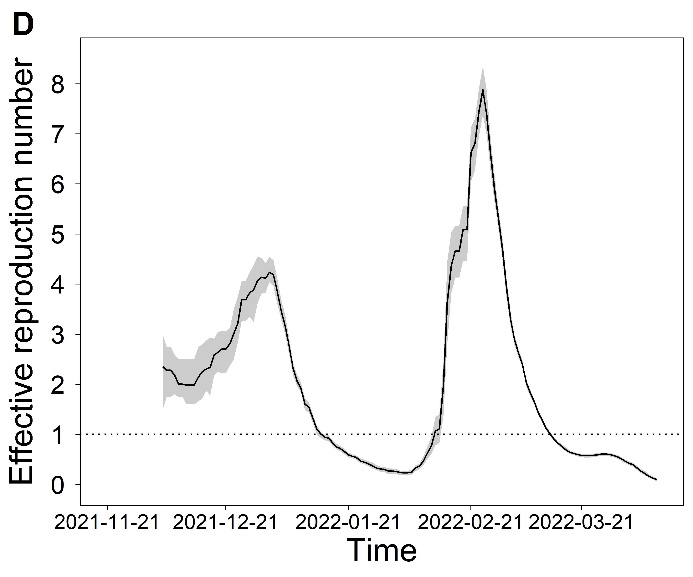

**Figure S2. Sensitivity of the effective reproduction number to the choice of serial interval distribution. A, B: discrete distributions of the serial interval of the model with the highest R-squared value. C, D: effective reproduction number estimated with these serial interval distributions. A, C: 2020-21 (H5N8), serial interval distribution with mean 2.72 days and standard deviation 1.07 days; B, D: 2021-22 (H5N1), serial interval distribution with mean 11.25 days and standard deviation 2.19 days.**
